# Brachiopod and mollusc biomineralisation is a conserved process that was lost in the phoronid-bryozoan stem lineage

**DOI:** 10.1101/2022.07.01.498437

**Authors:** Joel Vikberg Wernström, Ludwik Gąsiorowski, Andreas Hejnol

## Abstract

**Background:** Brachiopods and molluscs are lophotrochozoans with hard external shells which are often believed to have evolved convergently. While palaeontological data indicates that both groups are descended from biomineralising Cambrian ancestors, the closest relatives of brachiopods – phoronids and bryozoans – are mineralised to a much lower extent and are comparatively poorly represented in the Palaeozoic fossil record. Although brachiopod and mollusc shells are structurally analogous, genomic and proteomic evidence indicates that their formation involves a complement of conserved, orthologous genes. Here, we study a set of genes comprised of three homeodomain transcription factors, one signalling molecule and 6 structural proteins which are implicated in mollusc and brachiopod shell formation, search for their orthologs in transcriptomes or genomes of brachiopods, phoronids and bryozoans, and present expression patterns of 8 of the genes in postmetamorphic juveniles of the rhynchonelliform brachiopod *Terebratalia transversa*.

**Results:** Transcriptome and genome searches for the 10 target genes in the brachiopods *T. transversa, Lingula anatina, Novocrania anomala*, the bryozoans *Bugula neritina* and *Membranipora membranacea*, and the phoronids *Phoronis australis* and *Phoronopsis harmeri* resulted in the recovery of orthologs of the majority of the genes in all taxa. While the full complement of genes was present in all brachiopods with a single exception in *L. anatina*, a bloc of four genes could consistently not be retrieved from bryozoans and phoronids. The genes *engrailed, distal-less, ferritin, perlucin, sp1* and *sp2* were shown to be expressed in the biomineralising mantle margin of *T. transversa* juveniles.

**Conclusions:** The gene expression patterns we recovered indicate that while mineralised shells in brachiopods and molluscs are structurally analogous, their formation builds on a homologous process that involves a conserved complement of orthologous genes. Losses of some of the genes related to biomineralisation in bryozoans and phoronids indicate that loss of the capacity to form mineralised structures occurred already in the phoronid-bryozoan stem group and supports the idea that mineralised skeletons evolved secondarily in some of the bryozoan subclades.

## Background

Biomineralisation was an evolutionary innovation of major importance to early animals [1,2], its significance evident both from the vast amounts of skeletal taxa preserved as fossils and the ubiquity of hard tissues in present ecosystems. Beginning with a surge of small shelly fossils in terminal Ediacaran strata [3,4] the evolution of skeletons accelerated during the Cambrian radiations [5–8], and today a wide array of mineralised structures of differing chemical compositions are distributed across animal phylogeny. Many fascinating configurations of such hard parts can be found in the Lophotrochozoa, a subclade of Spiralia which includes brachiopods, annelids, molluscs, and several other invertebrate groups [9–14]. By the Early Cambrian; most lophotrochozoan crown groups had diverged and some had evolved biomineralised parts [4,7,8,15–17], leaving behind a trail of skeletal fossils [3,7,8] from an evolutionary arms race which culminated in the emergence of extensively mineralised taxa such as brachiopods and molluscs [17–21].While brachiopod shells are superficially similar to the shells of bivalve molluscs, molecular data [12,22–24] do not support the idea that brachiopods and molluscs have a close relationship within Lophotrochozoa. Rather, the favoured view is that the morphological similarities between brachiopod and mollusc shells are an example of homoplasy, the structures having arisen independently in an instance of convergent evolution [8]. In fact, even the shells and sclerites of different mollusc classes are considered likely to be independently derived structures [8,25–27] due to differences in form, formation, and composition. Within Lophotrochozoa, the closest relatives of brachiopods are instead the worm-like phoronids and the sessile bryozoans [28], and these three groups together form the clade Lophophorata [12,29–31]. Mineralised tissues are not evenly distributed among the lophophorates; no extant phoronids and only two out of five major clades of bryozoans are skeletal [32,33], while even the earliest known brachiopods had sclerites or shells [15,18–21]. Loss of biomineralised tissues seems to be a recurring evolutionary theme among lophotrochozoans (Fig. 1A): the putative phoronid stem group was mineralised [8,15,34,35] although extant phoronids are not, and similar losses seem to have occurred in annelids [36] and entoprocts [37]. Three clades of brachiopods survive to the present; the Rhynchonelliformea, Craniiformea and Linguliformea, with the latter two being most closely related [38,39]. In the extant rhynchonelliform brachiopod *Terebratalia transversa* Sowerby 1846, shell formation begins shortly after its three-lobed lecithotrophic larva has settled on appropriate substrate and given rise to a juvenile which resembles a miniature version of the adult animal [40]. After metamorphosis, a thin, unmineralised and non-articulating bivalved shell known as the protegulum envelops the juvenile [40]. Mineralisation of the protegulum begins shortly after metamorphosis is completed and takes place periodically [40] along the anterior and lateral sides of the structure. At the edge of the mantle epithelium (Fig. 1B) lies the periostracal slot where the shell is secreted. Lobate cells located beneath the periostracal slot produce the organic layer of the periostracum, while vesicular cells above them secrete calcite crystals in what is known as the ‘primary mineralisation’ [41,42]. Combined with shell material produced by the outer mantle epithelium in a ‘secondary mineralisation’, this eventually results in a calcareous shell marked by concentric growth lines [41,42], which articulates at a teeth- and-socket hinge structure [41].

**Figure 1.**
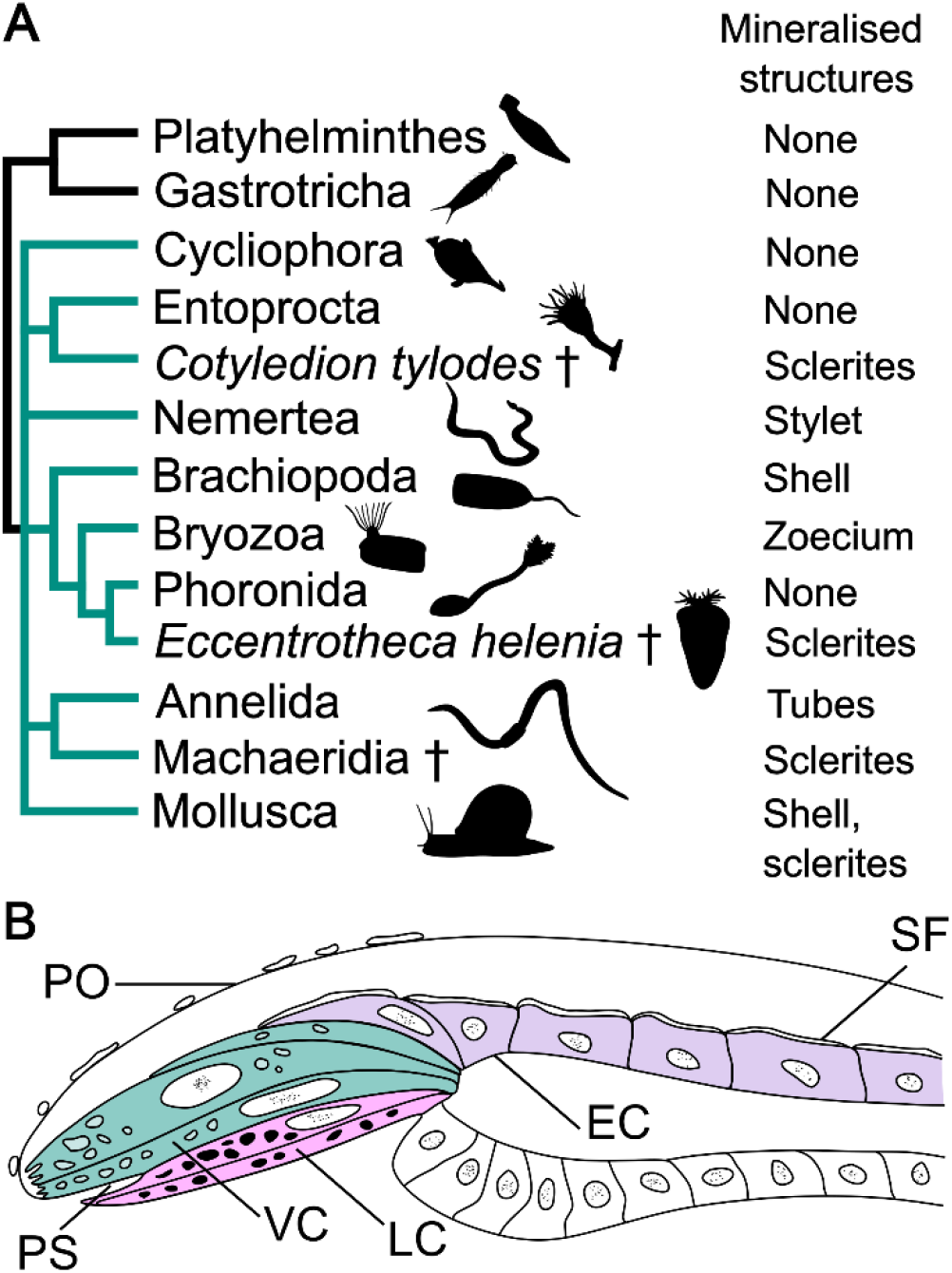
A. Cladogram depicting lophotrochozoan interrelationships [23,43,44] (in red) with Rouphozoa [45] as outgroup (in black) with an overview of mineralisation capacities. Taxa marked with an obelisk are Cambrian stem groups of their respective sister clades. B. Detailed morphology of the mantle margin which excretes the shell of the brachial valve. Lobate cells (pink) secrete the organic portion of the periostracum while vesicular cells (green) produce the primary layer of the inorganic shell and the outer epithelial cells of the mantle (purple) secrete the secondary layer of the inorganic shell. EC: outer mantle epithelial cell, LC: lobate cell, PO: periostracum, PS: periostracal slot, SF: fibre of the secondary shell layer, VC: vesicular cell.

By merit of being the best-studied extensively mineralised lophotrochozoan taxon, molluscs provide a useful reference point for the investigation of hard tissue formation in brachiopods. While the last common ancestor of the two groups was likely weakly mineralised at most [8], mollusc shell formation does have similarities to that of brachiopods [40,42,47,48], and mounting evidence suggests that the two groups may share some of the underlying genetic architecture required for biomineralisation [48–53]. A conserved ‘biomineralisation toolkit’ has been proposed by several authors [1,8,27,54–59], possibly encompassing not only molluscs, brachiopods and other lophotrochozoans but all bilaterians. This idea implies that while structurally analogous, the formation of shells in brachiopods and molluscs builds on a complement of orthologous genes, and has support from genomics- [49,51,57] and proteomics-based studies [53]. Recent single-cell transcriptomic data from the bivalve *Dreissena* have supported the notion of a biomineralisation toolkit but also showed enrichment of numerous lineage specific transcripts in the shell field, indicating rapid evolution of shell matrix proteins in molluscs [56]. Here, we have examined which of a handful of select mollusc genes related to biomineralisation are present in the transcriptomes or genomes of various lophophorate taxa and studied the expression patterns of the same genes in postmetamorphic juveniles of *T. transversa*. Our primary aims were to answer whether phoronids and non-mineralised bryozoans could have lost parts of their biomineralisation-related gene complement, and whether the genes would be expressed in the *T. transversa* mantle margin, where biomineralisation occurs [40–42]. We targeted 10 genes with a known function in the biomineralisation of molluscs and/or brachiopods, including the three homeodomain transcription factors *engrailed* (*en*) [48,56,59–65], *distal-less* (*dlx*) [66,67], *goosecoid* (*gsc*) [68], the *decapentaplegic* ortholog *bmp2-4* [59,64,66,69,70], the matrix protein genes *ferritin* [66,71–73], *calmodulin* [50] and *perlucin* [50,74,75], a mantle peroxidase [48,76,77], and two supposedly brachiopod-specific shell-associated genes [53] originally designated as F10023803 and R20087389 but herein dubbed shell protein 1 and 2 (abbreviated as *sp1* and *sp2*).

## Results

### The lophophorate biomineralisation gene complement and gene orthology

Our search for biomineralization-related genes in the transcriptomes and genomes of lophophorates resulted in the identification of several putative orthologs, the homology of which were tested using phylogenetic analyses (Add. Figs. 1-6). Brachiopods have retained orthologs of nearly all the targeted biomineralisation genes (Fig. 2), with the exception for an apparent loss of the specific *calmodulin* orthologue in *L. anatina*. Most of the genes are also identifiable in transcriptome or genome sequences from phoronids and bryozoans, with the notable exception of *perlucin* and *mpox*, in addition to the putatively brachiopod-specific genes *sp1* and *sp2*. Although sequences with similarity to *en* were present in the bryozoan transcriptomes, they did not form a clade with *en* sequences from other taxa (Add. Fig. 1). The seeming absence of some genes here signifies lack of an ortholog to the reference sequence, and does not indicate definite absence of other, closely related genes (for instance, several *calmodulin*-related genes are present in the *L. anatina* genome [49], but the specific *calmodulin* ortholog investigated here is not). Additionally, since we did not have access to genomes for all taxa and therefore only searched the transcriptomes of *B. neritina, M. membranacea, N. anomala* and *P. harmeri*, it is possible that some genes are present in the genome but were not expressed at the time of transcriptome generation.

**Figure 2.**
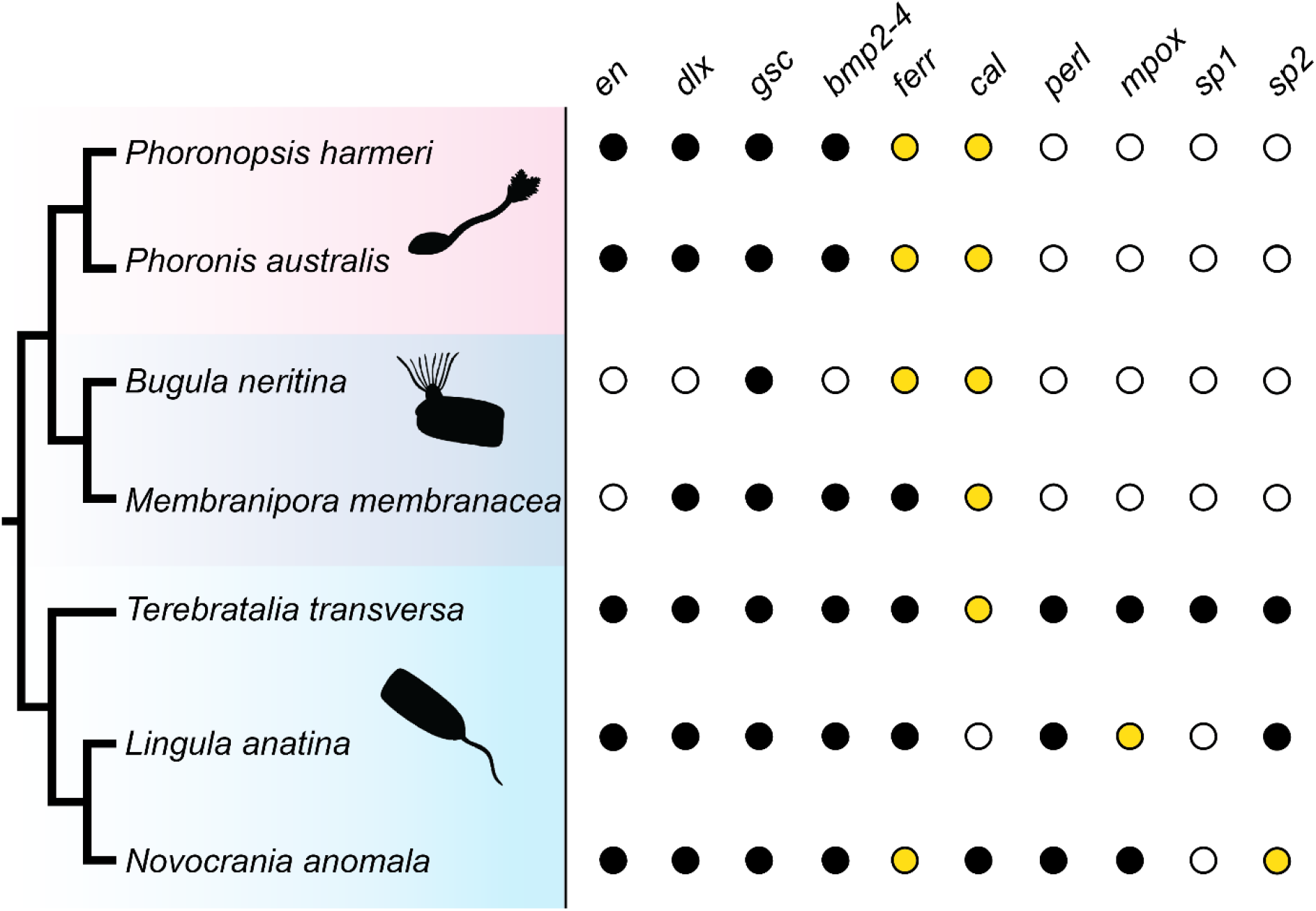
Cladogram depicting lophophorate relationships and presence-absence matrix of genes for each taxon. Filled dots indicate that an ortholog was found in the available genome/transcriptome, hollow dots indicate the opposite. Yellow dots indicate that multiple orthologs were identified.

### Expression patterns of biomineralisation genes in *T. transversa*

In-situ hybridisation of the genes yielded a range of expression patterns in various morphological features (Fig. 3A) of 2- and 6-day old juveniles of *T. transversa*, with most being expressed in varying degrees in the mantle margin, around the pedicle opening or in the area of the future hinges of the shell. Of the three homeodomain transcription factors, *en* was expressed exclusively in the mantle margin (black arrowhead, Fig. 3B). *dlx* was expressed in the mantle margin (black arrowheads, Figs. 3D and E) and lophophore (red arrowheads, Figs. 3D and E), while *gsc* was exclusively expressed in the lophophore (red arrowheads, Figs. 3F and G). The primary expression domains of *bmp2-4* were in the developing hinges of the protegulum (white arrowheads, Figs. 3H and I) and the lophophore (red arrowheads, Figs. 3H and I). Additionally, in juveniles at 6 dpm there was expression in the area of the future pedicle opening (blue arrowhead, Fig. 3I). *ferritin* was universally and abundantly expressed throughout the juvenile tissues to the point where overstaining became apparent minutes after starting the colour-producing reaction but seemed particularly prominent in the mantle margin (black arrowhead, Fig. 3C) and in an area posterior to the lophophore. *perlucin, sp1* and *sp2* were primarily expressed in the mantle margin (black arrowheads, Figs. 3J-O), with an additional *sp1* expression domain located in the outer epithelial cells of the mantle in juveniles at 2 dpm (yellow arrowhead and insert, Fig. 3L).

**Figure 3.**
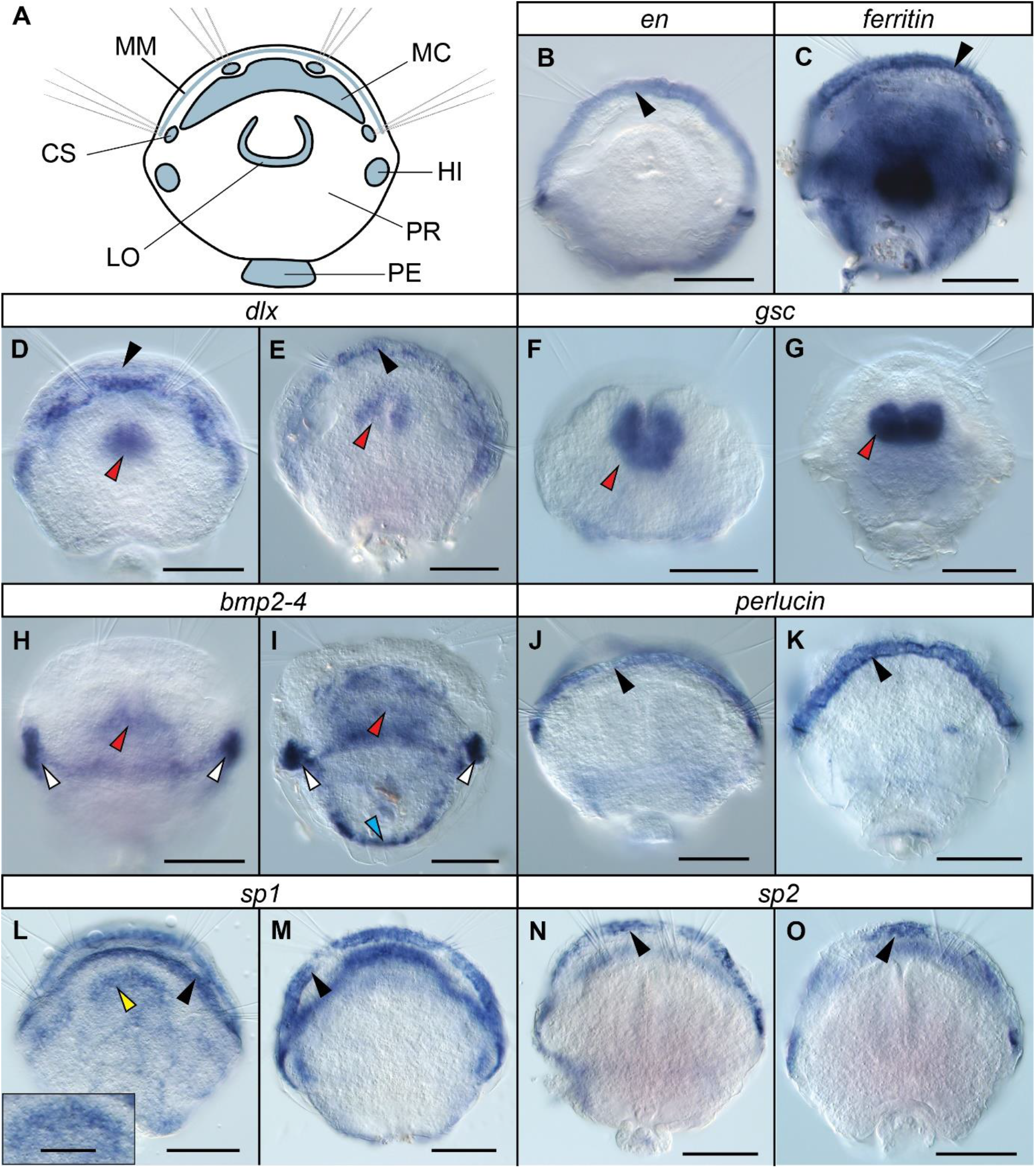
A. Morphological features of a *T. transversa* juvenile at 2 dpm, interpreted after Stricker & Reed [40] and Gąsiorowski & Hejnol [46]. B-O Expression patterns of targeted genes. Gene names are indicated in the white bars above the images. For each gene, the left picture depicts a juvenile at 2 dpm, and the right picture a juvenile at 6 dpm. Data could not be retrieved for *en* and *ferritin* at 2 dpm. The insert in panel L provides a magnified view of expression within the outer mantle epithelium cells. Black arrowheads indicate expression in the mantle margin, yellow arrowheads in the mantle epithelium, red arrowheads expression in the lophophore, white arrowheads expression in the hinges and blue arrowheads expression in the pedicle opening. Scale bars represent 50 μm (insert in panel L 20 μm). CS: chaetal sac, LO: lophophore rudiment, MC: mantle cavity, MM: mantle margin, PE: pedicle, PR: protegulum, HI: future hinge.

## Discussion

### A shared brachiopod-mollusc complement of genes involved in biomineralisation

The gene expression patterns we observed indicate that the two main groups of lophotrochozoan biomineralisers, brachiopods and molluscs, base their shell formation on a shared and conserved genetic foundation. Of the studied genes, *en* is arguably the most implicated in shell development in both groups [48,66] and as demonstrated here *en* was clearly and exclusively expressed in the *T. transversa* mantle margin, which indicates its involvement in shell formation due to the vicinity of CaCO_3_-excreting vesicular cells. *dlx* was also predominately expressed in the mantle margin, with an additional expression domain in the lophophore rudiment. Although *dlx* could not be confirmed to have biomineralising functions in *Lingula anatina* [48], the expression pattern herein corresponds well with the reported involvement of this gene in gastropod shell formation [66]. In accordance with our expression patterns in *T. transversa, dlx* is involved in lophophore formation in *L. anatina* [48], and the gene has been reported to be part of an “head appendage” genetic program [78] which agrees well with the idea that lophophores are homologous to the head of other bilaterians. Meanwhile, *gsc* was exclusively expressed in the developing lophophore and does not seem involved in *T. transversa* biomineralisation. Aside from its role in gastropod shell field formation, the gene is generally known to be one of the head patterning’ homeodomain transcription factors [79,80] with a specific role of patterning sensory cells around the mouth, agreeing well with the aforementioned link between lophophores and bilaterian heads [38,46,49,81]. *bmp2-4* was primarily expressed in the region of the lophophore, the developing hinge structure and the pedicle opening, the two latter of which are not close to the mantle margin but are areas which undergo structural modifications involving secondary biomineralisation of the pre-existing larval protegulum [40,42]. The *bmp2-4* ortholog in *Saccostrea kegaki* is likewise expressed in the hinges of the shell and seemingly important for hinge formation [64], and this similar expression pattern could represent a convergently evolved co-option of the gene for hinge formation. Lophophore expression of *bmp2-4* in the lophophore agrees with another proposed function of this gene family; the formation of the head, appendages and other outgrowths of the body [78,82]. The two shell matrix protein genes *ferritin* and *perlucin* were united by a clear expression in the mantle margin, but while this was the exclusive expression domain for perlucin, *ferritin* was broadly expressed also in other tissues. The highly specific mantle margin expression of *perlucin* adds weight to the argument for a shared brachiopod-mollusc biomineralisation gene complement, while the broad expression of *ferritin* could be explained by the wide range of physiological functions ferritins are known to carry [83]. Orthologues of *sp1* and *sp2*, the genes which were previously described as shell forming proteins from the rhynchonelliformean brachiopod *Magellania* [53] are also present in the transcriptomes or genomes of *Terebratalia, Novocrania* and *Lingula* but are absent in the transcriptomes or genomes of phoronids and bryozoans, indicating that the genes are brachiopod-specific and evolved in the common ancestor of all brachiopods after the split from other lophophorates. Alignment of the protein sequences of the *sp1* and *sp2* genes (Add. Fig. 7) suggests that both proteins are homologous, however they are divergently distributed among particular brachiopod clades. In *Lingula, sp2* was present but *sp1* absent, and in *Novocrania* we identified two paralogous sequences corresponding to *sp2*. Like in *Magellania*, both *sp1* and *sp2* could be identified in *Terebratalia*, suggesting that *sp1* may be a rhynchonelliform innovation. Both *sp1* and *sp2* were distinguished by mantle margin expression in *T. transversa* juveniles, indicating a conserved function among brachiopods. The expression of *sp1* in the outer epithelial cells of the mantle in 2-but not 6-dpm juveniles of *T. transversa* corresponds well with what would be expected of a gene involved in the progressive secondary mineralisation of the protegulum, which is finished shortly after metamorphosis [40]. In a broader context, our results support the notion of structurally analogous but genetically homologous shells in brachiopods and molluscs, an idea reinforced by both genomics- [49,51,57] and proteomics- [53] based studies. In addition to the genes investigated here, *pou3*, which regulates expression of shell matrix proteins in molluscs [54,84] is also expressed in the mantle margin of *T. transversa* [85], further supporting idea of a common molecular control of the biomineralisation in both groups. The fossil record likewise indicates that both molluscs [7,8,17,86] and brachiopods [15,18,19,21,87] have had the capacity for biomineralisation for a very long time, although their stem groups bore variations of sclerites rather than fully formed shells. Likely, their common ancestor had a genetic capacity for biomineralisation which was initially used to mineralise an organic skeleton [8] before giving rise to *bona fide* brachiopod and mollusc shells through convergent evolution.

### Implications for the evolution of lophophorate biomineralisation

It is not surprising that of the lophophorate clades, phoronids and bryozoans seemingly retain the fewest orthologs of the putative biomineralisation-related genes (Fig 2). It is worth noting that four of the tested genes which are present in the phoronid and bryozoan genomes or transcriptomes (*en, dlx, gsc* and *dmp2-4*) are not strictly biomineralisation-specific, but play multiple functions during the development of various lophophorates as has been shown in several studies [48,88–90]. When it comes to the more biomineralisation-specific genes, phoronids and bryozoans are united not only by the lack of tentatively brachiopod-specific *sp1* and *sp2* genes, but also of perlucin and mantle peroxidase which otherwise have orthologs in other bilaterians, including molluscs (Add. Figs. 5 & 6). Additionally, phoronids and bryozoans both lack the *Hox* gene *scr* [38,81,91], which in brachiopods is specifically expressed in the shell forming epithelium [46,49,92]. The loss of *scr* has previously been hypothesised to be related with the reduction of the biomineralisation capacities in phoronids [38,81]. The similar pattern of loss in bryozoans and phoronids agrees well with recent studies that indicate a sister-group relationship between the two [31,43,44,93,94]. The bryozoan stem group was unmineralised and originated in the early Cambrian, as indicated by recent fossil discoveries [95], while the crown group did not appear until at least in the Early Ordovician by which time the Stenolaemata had secondarily evolved a mineralised skeleton [96]. The notion of an unmineralised origin of bryozoans has further support [33], with the morphologically disparate skeletons of stenolaemates and cheilostomates likely being secondarily evolved as indicated by their phylogenetic distribution (Fig. 4). The sister relationship and similar pattern of loss of biomineralisation genes in phoronids and bryozoans suggests that the capacity of forming hard tissues was lost before the two groups diverged from each other. Taking this scenario into account, we suggest that the Early Cambrian tommottiid *Eccentrotheca helenia*, a putative stem group phoronid [34,35] which possessed a hard skeleton of loosely associated sclerites, likely represents an evolutionary stage predating loss of biomineralisation and should be rather considered as belonging to the phoronid-bryozoan stem group (Fig. 4).

**Figure 4.**
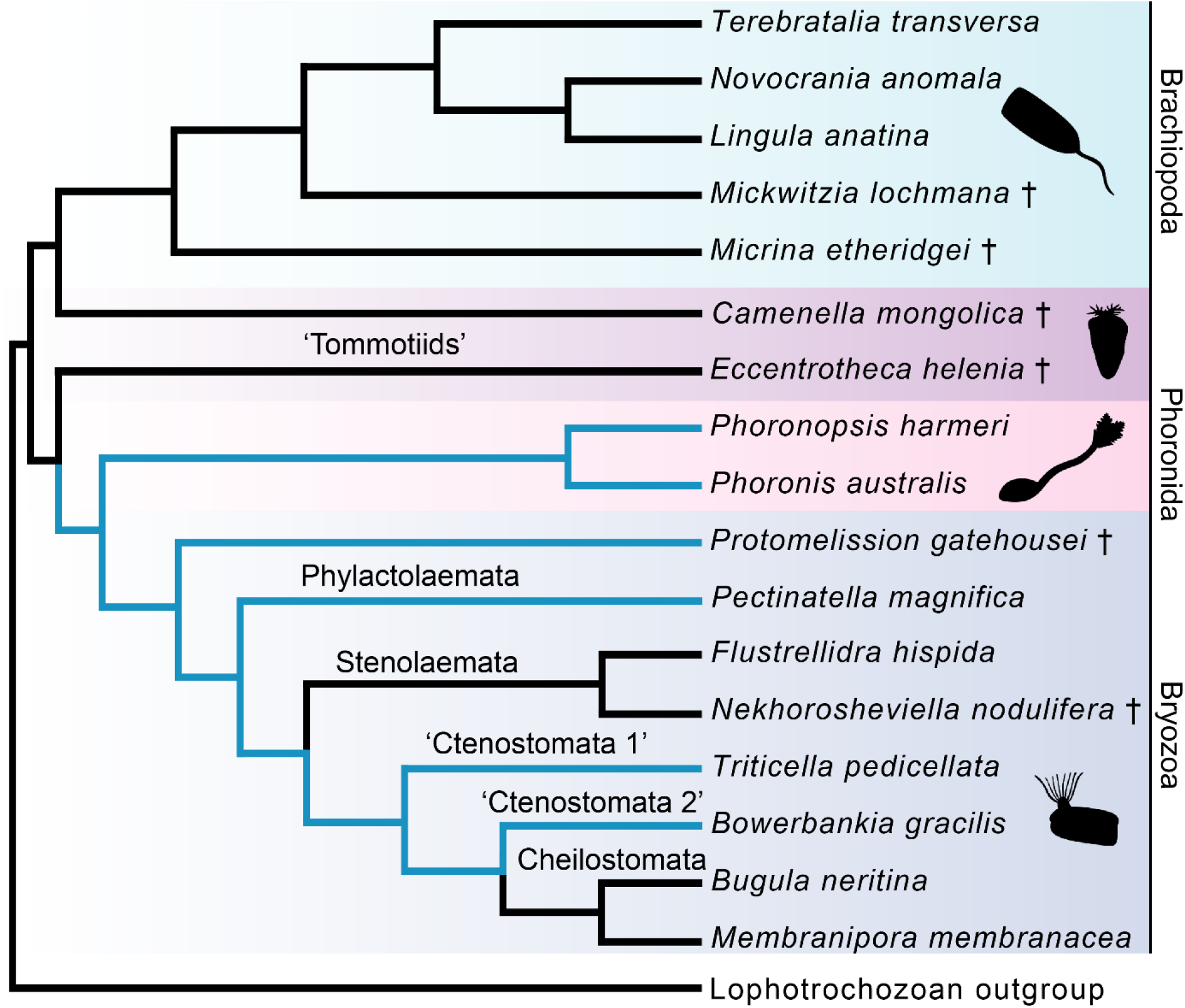
The evolution of biomineralisation in lophophorates. Black parts of the tree indicate biomineralising lineages, while light blue indicates soft-bodied taxa. The ancestral lophophorate has been assumed to be biomineralising due to the similarity between brachiopod and mollusc biomineralisation-related genes. Bryozoan topology after Schwaha *et al*. [33], Xia *et al*. [96] and Zhang *et al* (Zhang et al., 2021).

## Conclusions

Several genes involved in mollusc biomineralisation are expressed in the mantle margin and other biomineralising cell types in postmetamorphic juveniles of the rhynchonelliform brachiopod *Terebratalia transversa*. We suggest that the common ancestor of brachiopods and molluscs was able to form mineralised structures, for which it utilised a molecular machinery that has been at least partially retained in both lineages. This ancestral biomineralisation ability has been independently co-opted into formation of the shell-like external skeleton in molluscs and brachiopods, which evolved convergently as indicated by comparative morphology and the fossil record. This explains why the shells of both groups, despite being structurally analogous, show homology on the genetic level. Losses of some of the biomineralization-specific genes in bryozoans and phoronids indicate loss of the capacity to form mineralised structures in the phoronid-bryozoan stem group and supports the idea that mineralised skeletons evolved secondarily in some of the bryozoan subclades.

## Methods

### Specimen collection and preservation

Egg-bearing specimens of the rhynchonelliform brachiopod *Terebratalia transversa* were collected off San Juan Island, Washington State, USA. Animal husbandry and fertilisation of eggs was performed in vitro, and juveniles were fixed in 3.7 % formaldehyde and stored in methanol at -20 °C.

### Transcriptome searches and orthology assessment

A set of reference sequences for each gene of interest (Add. Table 1) was used to search the transcriptomes and genomes for orthologous sequences via BLAST. Transcriptomes of the bryozoans *Membranipora membranacea* and *Bugula neritina*, the brachiopods *Terebratalia transversa* and *Novocrania anomala* and the phoronid *Phoronopsis harmeri* (available in-house) and published genomes of the brachiopod *Lingula anatina* (GenBank accession number: LFEI00000000.2) and the phoronid *Phoronis australis* (GenBank accession number: NMRA01000001.1) were also screened for orthologs using the reference sequences. Sequences retrieved from the transcriptomes and genomes were imported into the software CLC Main Workbench v. 7, translated to protein sequences, and aligned with related sequences from other taxa retrieved from the NCBI repository of protein sequences and other databases. Alignments of protein sequences (including the *sp1* and *sp2* genes, Add. Fig 7) were trimmed using the software trimAl v1.1 [97] and imported into FastTree v. 2.1.10 [98] for phylogenetic reconstructions using the ML Model option ‘Le-Gascuel 2008’ and CAT approximation with 20 rate categories. The resulting approximately-maximum-likelihood phylogenies (Add. Figs. 1-6) were used to assess orthology of the lophophorate genes of interest. New sequences obtained and identified in this study were uploaded to GenBank (accession numbers ON868422 to ON868458).

### Gene amplification, cloning and sequencing

Specific primers against the genes of interest were designed using the software MacVector v 11.1.2. Genes were amplified from cDNA libraries using PCR, verified by electrophoresis on a 1 % agarose gel, extracted from the gel using a MinElute Gel Extraction Kit according to the manufacturer’s instructions, and ligated into vectors. The vectors were heat-shocked into competent *Escherichia coli* cells which were left to proliferate for 24 hours in 37 °C. Bacterially amplified vectors were purified using a Qiagen Spin Miniprep Kit according to the manufacturer’s instructions, and inserts were sequenced using the Sanger method.

### Riboprobe synthesis, whole-mount in situ hybridisation and imaging

PCR was used to amplify genomic DNA from vectors, and amplicons were purified using a MinElute PCR Purification Kit according to the manufacturer’s instructions. Resulting nucleic acid concentrations were measured using a NanoDrop device, while amplicon sizes were verified through electrophoresis on a 1 % agarose gel. dUTP-digoxigenin-labelled RNA probes were synthesised according to the manufacturer’s instructions (Roche, USA). WISH was performed in accordance with previously published protocols [46,90,99]. Following specimen digestion with proteinase K for 10 minutes, probes were allowed to hybridise with *T. transversa* juveniles at a concentration of 1 ng ⋅μL^-1^ at 67 °C for 72 hours. Anti-digoxigenin-AP antibodies were used to detect the probes, and they were visualised using nitroblue tetrazolium chloride and 5-bromo-4-chloro-3-indolyl phosphate. Successfully hybridised and visualised specimens were transferred to cover slides, mounted in 70% glycerol and photographed using a Zeiss Axiocam camera connected to a Zeiss Axioscope Ax10 using Nomarski bright field optics.

## Declarations

### Ethics approval and consent to participate

Studies of brachiopods do not require ethics approval or consent to participate.

### Consent for publication

Not applicable.

### Availability of data and materials

The dataset of genetic sequences supporting the conclusions of this article is available in the GenBank repository (https://www.ncbi.nlm.nih.gov/genbank/) under accession numbers ON868422 to ON868458. All remaining data generated or analysed during this study are included in the article itself or its supplementary materials.

### Authors’ contributions

LG and JVW conceived the study, JVW conducted gene search and orthology assessments, performed in situ hybridisation, produced figures, and drafted the manuscript. LG, JVW and AH analysed data, and read, discussed, and approved the final manuscript.

## Acknowledgements

We are thankful to Ferenc Kagan for his input on some bioinformatic issues, and to Aina Børve for providing valuable assistance in the lab. The staff of UW Friday Harbor Laboratories and crew of the vessel “Centennial” are gratefully acknowledged for helping in collection of adult *T. transversa*. We would also like to thank all present and former members of the Hejnol lab who contributed to culturing and fixing *T. transversa* juveniles.

