## Additional Material for "Brachiopod and mollusc biomineralisation is a conserved process that was lost in the phoronid-bryozoan stem lineage"

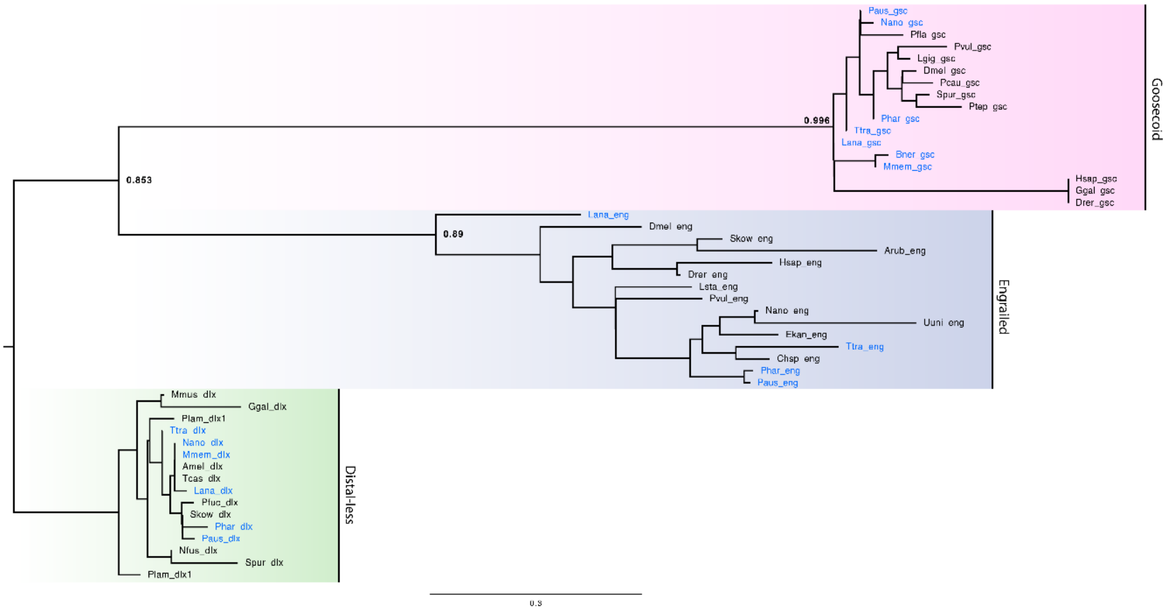


**Additional figure 1.** Phylogenetic relationships of the homeodomain transcription factor proteins engrailed, goosecoid and distal-less in various metazoan taxa. Blue font denotes lophophorates. SH-like support values are indicated for select nodes.

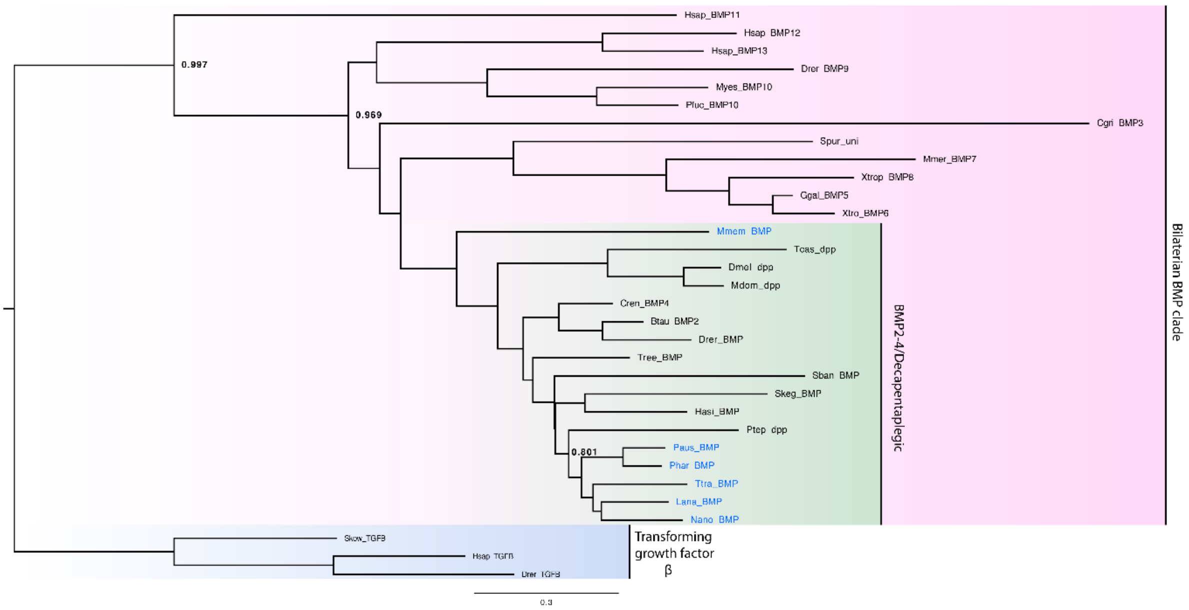


**Additional figure 2.** Phylogenetic relationships of the signalling molecule BMP2-4 to closely related proteins in various metazoan taxa. Blue font denotes lophophorates. SH-like support values are indicated for select nodes.

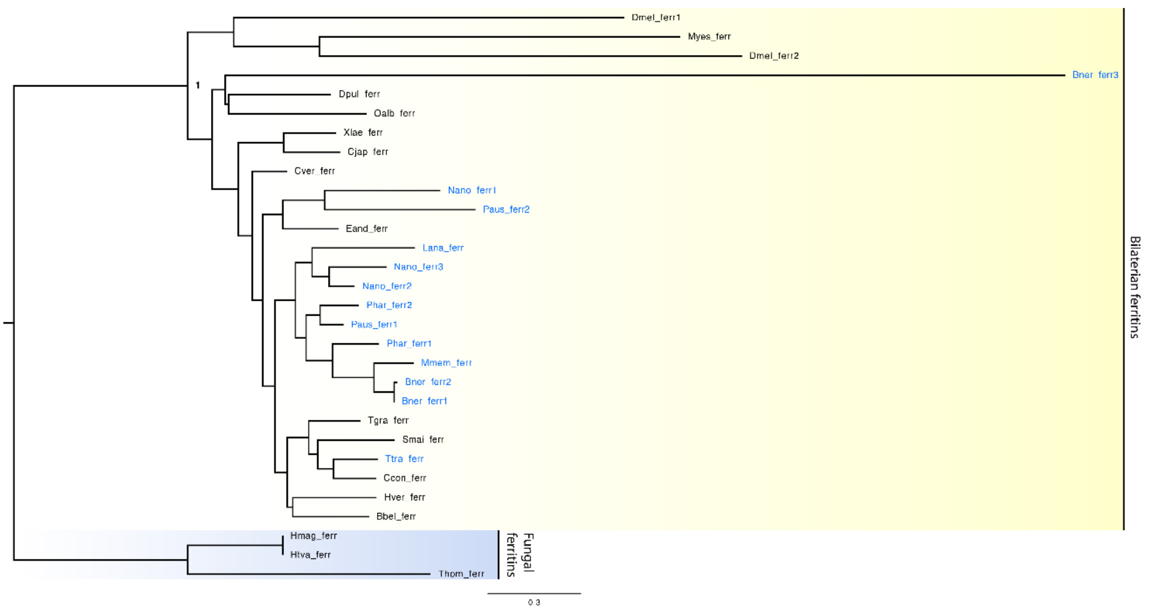


**Additional figure 3.** Phylogenetic relationships of ferritins from various metazoan taxa to fungal orthologues. Blue font denotes lophophorates. SH-like support values are indicated for select nodes.

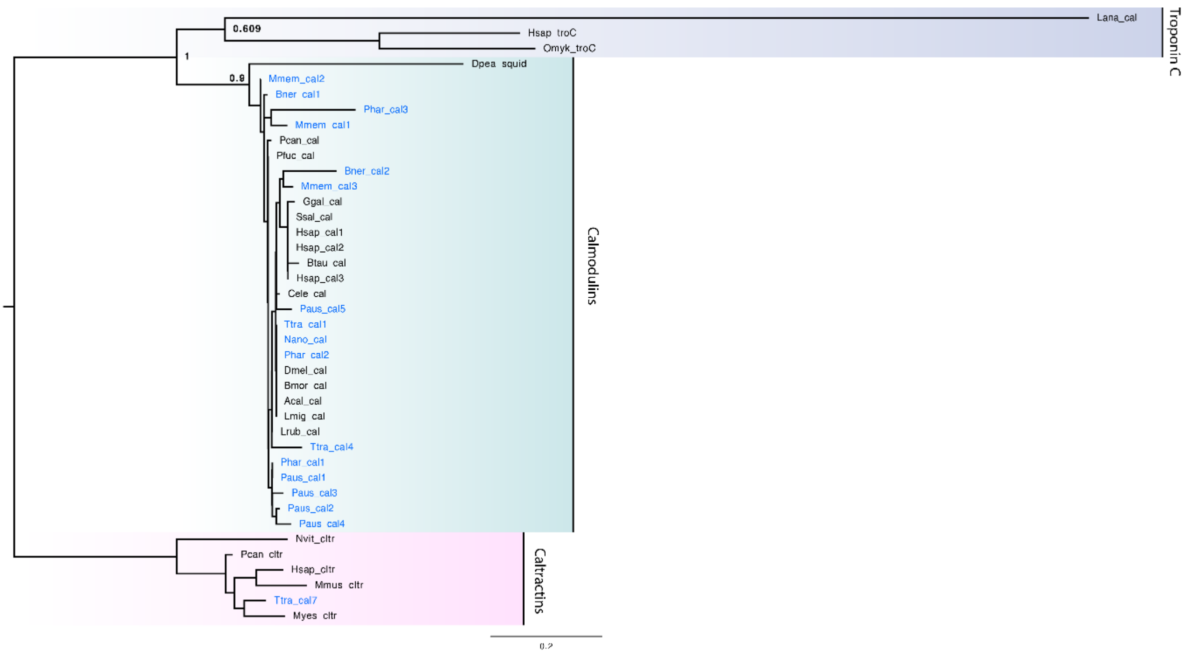


**Additional figure 4.** Phylogenetic relationships of calmodulin to other calcium-binding proteins in various metazoan taxa. Blue font denotes lophophorates. SH-like support values are indicated for select nodes.

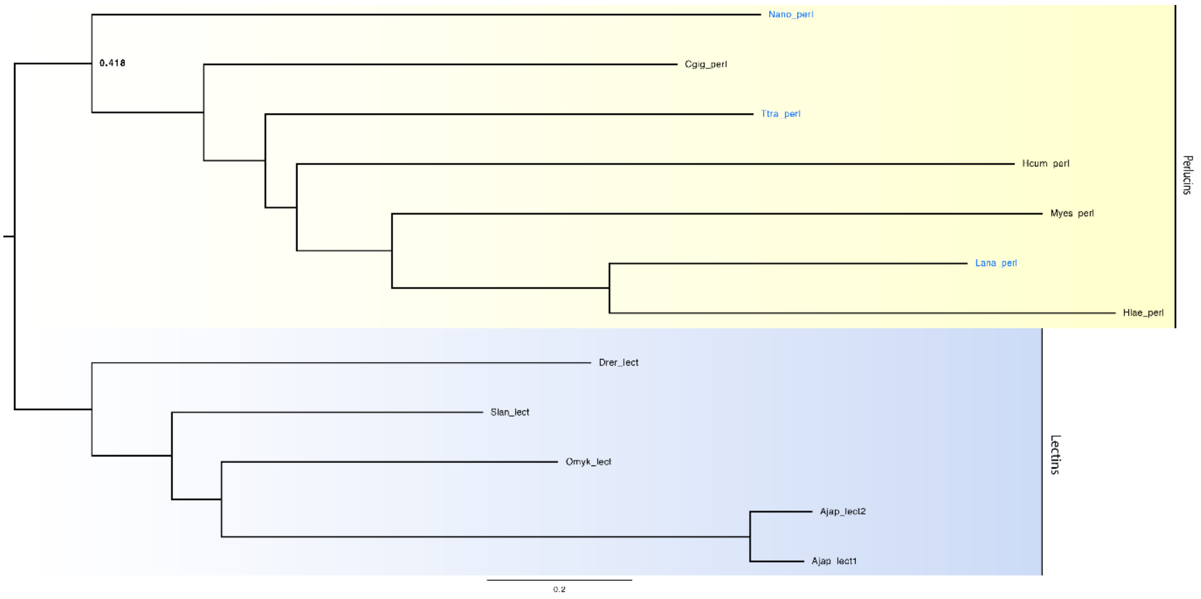


**Additional figure 5.** Phylogenetic relationships of perlucin orthologues to other lectin-related proteins in various metazoan taxa. Blue font denotes lophophorates. SH-like support values are indicated for select nodes.

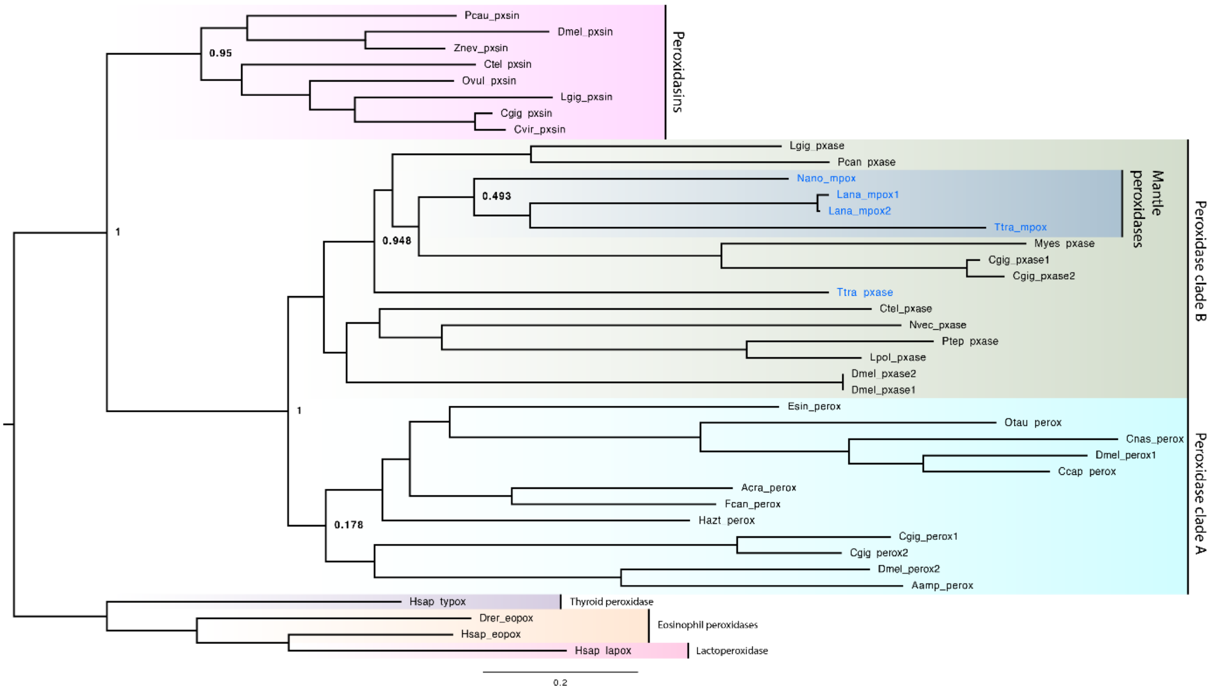


**Additional figure 6.** Phylogenetic relationships of mpox (mantle peroxidases) to other peroxidase-related proteins in various metazoan taxa. Blue font denotes lophophorates. SH-like support values are indicated for select nodes.

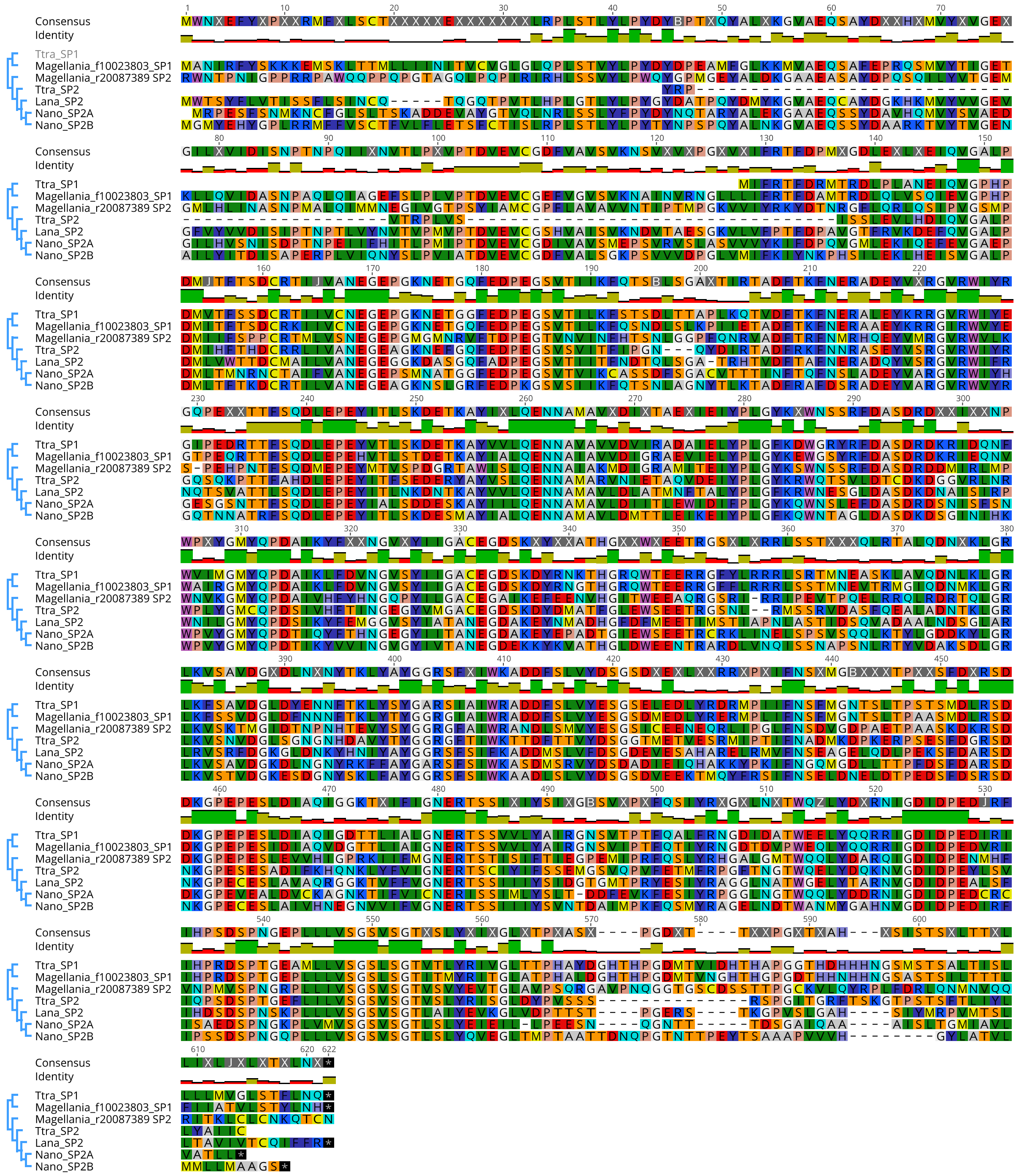

**Additional figure 7.** Alignment of the *SP1* and *SP2* protein sequences from four brachiopod species.

**Additional table 1.** Reference sequences used to search genomes/transcriptomes for each gene of interest.

| Gene name | Genbank accession | Organism | Notes |
|---|---|---|---|
| Engrailed | ALS19757.1 | *Terebratalia transversa* | |
| Distal-less | BBB21813.1 | *Lingula anatina* | |
| Decapentaplegic/BMP2-4 | XP_013396300.1 | *Lingula anatina* | |
| Goosecoid | XP_013379052 | *Lingula anatina* | |
| Mpox (mantle peroxidase) | BBB21814.1 | *Lingula anatina* | |
| Ferritin | XP_023933523 | *Lingula anatina* | |
| Calmodulin | NP_001191509.1 | *Aplysia californica* | |
| Perlucin | AGI61062.1 | *Hyriopsis cumingii* | |
| F10023803 (SP1) | | *Magellania venosa* | From Jackson et al. 2015 |
| R20087389 (SP2) | | *Magellania venosa* | From Jackson et al. 2015 |
